# Yeast-2-Hybrid-Seq and Bifluorescence Complementation Resources for assessing Protein:Protein Interactions in Arbuscular Mycorrhizal Roots: CKL2 as a Case Study

**DOI:** 10.1101/2025.08.07.669188

**Authors:** Sergey Ivanov, Lena M. Müller, François M. Lefèvre, Maria J. Harrison

## Abstract

Reverse genetics, facilitated by CRISPR technologies and comprehensive sequence-indexed insertion mutant collections, has advanced the identification of plants genes essential for arbuscular mycorrhizal (AM) symbiosis. However, a mutant phenotype alone is generally insufficient to reveal the specific role of the protein in AM symbiosis and in many cases, identifying interacting partner proteins is useful. To enable identification of protein:protein interactions during AM symbiosis, we established a *Medicago truncatula -Diversispora epigaea* yeast-two-hybrid (Y2H) library which, through Y2H-seq screening, can provide a rank-ordered list of candidate interactors of a protein of interest. We also developed a vector system to facilitate bimolecular fluorescence complementation assays (BIFC) in mycorrhizal roots so that protein interactions can be assessed in their native cell types and sub-cellular locations. We demonstrate the utility of a Y2H-seq screen coupled with BIFC in mycorrhizal roots, with a search for proteins that interact with CYCLIN DEPENDENT LIKE KINASE 2 (CKL2), a kinase essential for AM symbiosis. The Y2H-seq screen identified three 14-3-3 proteins as the highest ranked CKL2 interacting proteins. BIFC assays in mycorrhizal roots provided evidence for a CKL2:14-3-3 interaction at the periarbuscular membrane (PAM) in colonized root cells. Down-regulation of 14-3-3 by RNA interference provides initial evidence for a function in AM symbiosis. Thus, CKL2 may utilize 14-3-3 proteins to direct signaling from the PAM. The Y2H and BIFC resources will accelerate understanding of protein functions during AM symbiosis.

## I. Introduction

The arbuscular mycorrhizal symbiosis, a mutualistic association of plants and fungi from the Glomeromycotina, arose over 450 million years ago and has been retained by the majority of the plant families (Delaux & Schornack, 2021). Today it exists in terrestrial ecosystems across the planet, where it generally has a positive impact on plant health by enhancing mineral nutrient acquisition and also providing protection against pathogens (Sawers *et al*., 2018; Genre *et al*., 2020; Lutz *et al*., 2023)

AM symbioses are endosymbiotic associations and their development is complex with requirements for continual signaling and the coordinated differentiation of both root cells and fungal hyphae to generate the interfaces for reciprocal nutrient exchange (MacLean *et al*., 2017; Lanfranco *et al*., 2018; Wang *et al*., 2022). Under permissive conditions, generally phosphate deprivation, strigolactones and chitooligosaccharide-based molecules activate signaling in the fungus and plant, respectively. This signal activation underlies cellular changes that enable fungal hyphae to grow through the epidermal cells and into the root cortex (Genre & Russo, 2016; Zipfel & Oldroyd, 2017; Hull *et al*., 2021; Das *et al*., 2022). Here, the main endosymbiotic phase ensues and fungal hyphae differentiate within the root cortical cells, generating so-called arbuscules (branched hyphae) (Choi *et al*., 2018). Each arbuscule is surrounded by a newly developed plant membrane, the periarbuscular membrane (PAM), which contains transporters that enable nutrient exchange between the two symbionts but also proteins typically involved in signaling, including kinases of several types (Harrison & Ivanov, 2017; Wang *et al*., 2017; Roth *et al*., 2018; Ivanov & Harrison, 2024). While development of the arbuscule-cortical cell interface is complex, the arbuscule lives only for a few days before disassembly occurs. The plant cell appears to return to its former non-colonized state while the fungus develops new arbuscules as hyphal growth progress along the root (Bonfante-Fasolo, 1984; Gutjahr & Parniske, 2013).

Knowledge of the molecular basis of the association is advancing; the plant’s AM symbiosis-associated transcriptome, that underlies the metabolic, physiological and cellular modifications in the root cortical cells, has been well-documented in many plant species and was recently extended to the single cell level (Radhakrishnan *et al*., 2020; Carrere *et al*., 2021; Shi *et al*., 2021; Serrano *et al*., 2024; Sgroi *et al*., 2024). Forward and reverse genetic screens, the latter often guided by transcriptional data, have defined plant genes required for AM symbiosis and identified transcriptional regulators, metabolic pathways and cellular processes central to the association (reviewed in (MacLean *et al*., 2017; Delaux & Schornack, 2021; Wang *et al*., 2022). However, to gain a deeper understanding, it is essential to determine how these proteins function to facilitate symbiosis; in many cases, this requires knowledge of protein-protein interactions, particularly for the new signaling proteins whose partners and pathways are unknown. Analyses of protein:protein interactions in colonized cells is particular challenging as these cells comprise a small proportion of the root system and the dynamics of arbuscule development and degeneration adds further complexity (Serrano *et al*., 2024).

Recently, we considered these challenges as we initiated experiments to determine the functions of two cyclin-dependent like kinases, CKL1 and CKL2 (Ivanov & Harrison, 2024). These proteins are essential for AM symbiosis and are associated, via myristoylation, with the PAM and plasma membrane (PM) specifically in colonized root cortical cells. Membrane association of the CKL proteins is essential for their function and both CKL proteins interact with, and are phosphorylated by symbiosis receptor proteins, malectin domain leucine-rich repeat receptor-like kinase DMI2 and a subset of the LysM-domain receptor-like kinases. *ckl1 ckl2* mutants fail to express the WRINKLED transcriptional regulators which led to a model in which CKL signaling ultimately regulate aspects of lipid provisioning to the fungal symbiont (Ivanov & Harrison, 2024). However, CKL signaling pathway(s) are currently undefined.

Yeast-two-hybrid libraries generated from mycorrhizal root cDNA are not widely available. Here, we prepared a *Medicago truncatula/Diversipora epigaea* root Y2H library. We validated the library through a Y2H-seq screen (Erffelinck *et al*., 2018) with DOES NOT MAKE INFECTIONS 3 (DMI3), whose partner protein INTERACTING PROTEIN of DMI3 (IPD3) is known, and simultaneously, we screened for interactors of CKL2. The top CKL2 protein interaction candidates were then evaluated by BIFC assay in mycorrhizal roots with corresponding genes expressed from their native promoters. This relatively rapid and simple workflow allowed us to demonstrate that a member of the 14-3-3 protein family interacts with CKL2, predominantly at the PAM in *M. truncatula* colonized root cells.

## II. Materials and Methods

### 1. Plant and fungal materials

*Medicago truncatula* accession Jemalong A17 and AM fungi, *Rhizophagus irregularis* (strain 197198) (formerly *Glomus intraradices*) and *Diversispora epigaea* (formerly *Glomus versiforme)* were used in this study.

### 2. Growth conditions

*M. truncatula* plants or composite plants with transgenic roots were planted into 20.5 cm plastic cones filled with a sterile mixture of sand/gravel containing with 100 sterile spores of either *Rhizophagus irregularis* or *Diversispora epigaea* placed at a depth of 4 cm below the surface. Plants were grown in a growth chamber under a 16 hour light/22°C and 8 hour dark/20°C regime with 40% humidity. Half-strength Hoagland’s solution containing half-strength nitrogen and 50 μM potassium phosphate was supplied twice a week. Plants were harvested at 4-5 weeks post planting depending on the experiment.

### 3. Generation *of Medicago truncatula* composite plants

To obtain composite plants with transgenic roots, *A. rhizogenes* strain ARqual mediated root transformation was used (Boisson-Dernier *et al*., 2001). For sub-cellular localization or BiFC analysis transgenic roots were selected based on visual detection of fluorescent arbuscular mycorrhizal symbiosis specific markers carried on the T-DNA (*BCP1pro:NLS-2×mCherry*). For RNA interference, transgenic roots were selected based on visual detection of fluorescent root transformation marker carried on the T-DNA (RedRoot; *AtUBQ10pro:DsRed*).

### 4. *Medicago truncatula-Diversispora epigaea* cDNA library construction

The *Medicago truncatula-Diversispora epigaea (Mt-De) root* cDNA library was generated using the CloneMiner™ II cDNA Library Construction Kit from Invitrogen™. The *Mt-De* primary library consisted of 3.6 × 10^6^ CFU / µg mRNA with an average insert size of 1.1 kb. The library was transferred to pJG4-5 where proteins are expressed as fusions with a B42-activation domain for Y2H screens.

### 5. Yeast-2-hybrid sequencing (Y2H-seq) screen

The protocol followed that of Erffelinck et al., (Erffelinck *et al*., 2018) but used yeast strain EGY48 (mutated in the leucine biosynthesis gene LEU2) carrying the pSH818 plasmid (carrying the LacZ reporter gene) and Gateway-compatible vectors (pEG202, pJG4-5). While the Y2H-seq screen is based on selection on leucine dropout plates rather than LacZ, including the pSH818 plasmid ensures flexibility of the approach and allows additional validation of positive colonies on X-Gal media. Briefly, bait constructs containing *DMI3* or *CKL2* were generated as LexA-binding domain fusions in the yeast expression vector, pEG202, and transformed into EGY48-pSH818 following a high-efficiency yeast transformation protocol (Gietz & Schiestl, 2007). The resulting bait strains were transformed with the pJG4-5 empty vector and grown on leu-plates to assess autoactivation. For both DMI3 and CKL2, autoactivation was minimal. The *Mt-De* library in pJG4-5 was transformed into the DMI3 or CKL2 bait strains using the high efficiency yeast transformation protocol (Gietz & Schiestl, 2007). An initial plating on ura-his-trp-dropout media selected for transformants and these were collected and stored as an intermediate library. Following growth in raffinose and galactose liquid media to induce expression of prey proteins, a second plating onto selective media (ura-his-trp-leu-drop-out media) selected for interactors. A 2-3X coverage of the cDNA library requires screening ∼5×10^6^ CFU of the *Mt-De* library. Following growth on leu-plates, interaction positive colonies were washed from the plates and pooled, and plasmids were extracted using the Zymoprep II yeast mini prep kit. The inserts were amplified with primers B5940 and B2008 and amplicons purified using a Quiagen PCR purification. This procedure generated 1 technical replicate and was repeated twice resulting in three batches of amplicons per bait construct. Purified amplicons (1-2 µg) were sheared (Covaris sonicator E220) and used as input for Illumina libraries following the method of Zhong et al., (Zhong *et al*., 2011). The libraries were sequenced on an Illumina NextSeq 500 × 1 × 75bp (extended mode), 12 samples per lane. Illumina libraries were also prepared with amplicons generated from the JG4-5 *Mt-De* library DNA transformed into EGY48-pSH818 cells carrying the pEG202 empty vector (no bait) processed alongside the DMI3 and CKL2 selection strains to provide baseline knowledge of transcript abundance within the library. Three libraries were sequenced for each bait and for the library baseline control.

Reads were mapped to the *M. truncatula* genome sequence (Mt4), read counts determined and counts-per-million (cpm)-normalized counts for each bait protein were ranked based on abundance. The positive control screen with DMI3 returned the known interactor, IPD3/CYCLOPs (Yano *et al*., 2008; Horvath *et al*., 2011), as the top-ranked interacting protein (Table 1).

**Table 1.** Top-ranked interactors of DMI3 as identified in a Y2H-seq screen. Values shown are the average transcript counts (cpm) from three replicate libraries. The top 30 genes with highest enrichment in the DMI3-selected library are shown, alongside cpm values for the same genes in the CKL2-selected library and the baseline control in the *M. truncatula* – *D. epigaea* library (Mt-De).

|  | Gene ID | Annotation | average<br>*Mt-De* | average<br>DMI3 | average<br>CKL2 |
| --- | --- | --- | --- | --- | --- |
| 1 | Medtr5g026850 | IPD3/CYCLOPS | 28.90 | 965311.77 | 5.23 |
| 2 | Medtr7g090180 | S-adenosylmethionine-dependent methyltransferase (SAM7) | 45.70 | 293056.00 | 35.29 |
| 3 | Medtr2g080130 | S-adenosylmethionine-dependent methyltransferase, putative (SAM2) | 6.96 | 174525.30 | 1.31 |
| 4 | Medtr3g449200 | hypothetical protein | 27.56 | 61507.34 | 18.18 |
| 5 | Medtr4g035650 | chaperone dnaJ-like protein | 14.00 | 61014.59 | 7.52 |
| 6 | Medtr8g104210 | hypothetical protein | 15.99 | 59115.73 | 16.87 |
| 7 | Medtr7g009070 | thioredoxin H2-like protein | 35.60 | 58990.59 | 44.39 |
| 8 | Medtr1g054535 | carbon catabolite repressor-like protein | 3.73 | 54662.13 | 2.19 |
| 9 | Medtr4g070080 | RNA-binding family protein | 135.91 | 50947.86 | 159.82 |
| 10 | Medtr7g057340 | (E)-beta-caryophyllene synthase | 2.28 | 48794.80 | 0.00 |
| 11 | Medtr5g013240 | transmembrane protein, putative | 12.56 | 45064.85 | 27.08 |
| 12 | Medtr7g112860 | dormancy/auxin associated protein | 162.42 | 35369.49 | 264.60 |
| 13 | Medtr7g057390 | (E)-beta-caryophyllene synthase | 1.86 | 32789.88 | 0.36 |
| 14 | Medtr2g006170 | 60S ribosomal protein L27-1 | 40.05 | 32047.14 | 51.55 |
| 15 | Medtr4g116410 | 40S ribosomal protein S11-1 | 69.44 | 31821.25 | 75.15 |
| 16 | Medtr1g073940 | HR-like lesion-inducing protein | 22.24 | 31553.42 | 35.57 |
| 17 | Medtr1g071540 | kinase interacting (KIP1-like) family protein | 2.32 | 30312.81 | 1.30 |
| 18 | Medtr3g078130 | DSS1/SEM1 family protein | 31.12 | 29346.53 | 53.65 |
| 19 | Medtr4g021350 | aldo/keto reductase family oxidoreductase | 77.14 | 27020.56 | 50.42 |
| 20 | Medtr1g029140 | transmembrane protein, putative | 0.99 | 23519.15 | 0.74 |
| 21 | Medtr5g064500 | phospho-2-dehydro-3-deoxyheptonate aldolase | 33.02 | 22582.33 | 27.26 |
| 22 | Medtr4g071150 | core histone H2A/H2B/H3/H4 | 7.81 | 22514.23 | 244.98 |
| 23 | Medtr1g054545 | endonuclease/exonuclease/phosphatase family protein | 1.18 | 22242.11 | 0.39 |
| 24 | Medtr1g081410 | 40S ribosomal protein S24-2 | 200.26 | 21119.22 | 168.76 |
| 25 | Medtr2g089865 | wound-responsive family protein | 11.67 | 20329.51 | 21.62 |
| 26 | Medtr0147s00701 | 60S ribosomal protein L37-1 | 83.97 | 19790.83 | 116.11 |
| 27 | Medtr3g062450 | ubiquitin-conjugating enzyme E2 | 37.40 | 19490.89 | 44.58 |
| 28 | Medtr4g087100 | senescence-associated nodulin | 1.47 | 18797.08 | 0.27 |
| 29 | Medtr3g107717 | GTP-binding nuclear Ran-like protein | 86.23 | 18559.20 | 51.61 |
| 30 | Medtr3g09878 | Serine-threonine protein phosphatase 7 | 3.37 | 16486.12 | 2.05 |

### 6. Plasmid construction for confocal laser-scanning microscopy, BiFC assay and RNA interference analysis

*Medicago truncatula* accession Jemalong A17 cDNA was used as template for PCR amplification of coding sequences of 14-3-3A, 14-3-3B, STS, TMP3, WD, PHd, REM-CT, TMP2, and DMI3. *M. truncatula* genomic DNA was used as template for PCR amplification of promoter regions of STR1, STR2, 14-3-3A and DMI3 and gene genomic sequences of STR1, STR2 and 14-3-3A. *NLS-mCherry* was amplified using pENTRL1-L2 *NLS-mCherry-GUS* vector (Ivanov and Harrison, 2014) as a template. Primers, used for DNA amplification, are listed in Table S1. PCR fragments with flanking *att*B sites were recombined with either pDONR221 to obtain pENTR L1-L2 clones, pDONR P4P1R to obtain pENTR L4-R1 clones or pDONR P2RP3 to obtain pENTR R2-L3 using Gateway™ BP Clonase™ II Enzyme mix (ThermoFisher Scientific). The pENTR clones were used in three fragment recombination with destination vectors pK7m34GW or pK7m34GW-BCP-R (Ivanov and Harrison, 2024) using Gateway™ LR Clonase™ II Enzyme mix (ThermoFisher Scientific) to create expression vectors. pENTR clones and expression vectors created and used in this study are listed in Table S2. pENTR clones containing promoter regions, gene genomic or coding sequences of *CKL1* and *CKL2* were reported previously (Ivanov and Harrison, 2024).

#### Expression vectors for BiFC in Nicotiana benthamiana

To create expression vectors containing two expression cassettes of genes of interest in a single T-DNA for BiFC and co-immunoprecipitation analysis in *N. benthamiana*, the expression cassettes were amplified and cloned as described before (Ivanov & Harrison, 2024) and shown here in Fig. S1c). Briefly, expression cassettes for putative protein interactors of CKL2 were generated by Gateway cloning and then the cassette was amplified using B7952 and B7953 primers containing *Aar*I sites. The expression cassette *AtUBQ10pro:CKL2-cYFP* (or *AtUBQ10pro:CKL1-cYFP*) was amplified using B7954-B7955 primers containing *Aar*I sites (Table S1). The amplified cassettes were used in Golden Gate cloning reaction using the *Aar*I restriction enzyme (ThermoFisher Scientific) and T4 DNA Ligase (Promega) to create co-expression vectors for BiFC analysis in *N. benthamiana* (Table S3).

#### Expression vectors for BiFC in M. truncatula with a colonized cell-specific marker gene

To create a variant of pCAMBIA2300-AarI-*ccd*B (Ivanov & Harrison, 2024) which contains a nuclear located arbuscular mycorrhizal symbiosis specific marker, the expression cassette *MtBCP1pro:NLS-mCherry-mCherry-t35S (NLS-2×mCherry)* was amplified using primers B3901 and B4704, containing recognition sites for *Kpn*I and *Sac*I restriction endonucleases, and cloned at corresponding restriction sites of pCAMBIA2300-AarI-*ccd*B to create pCAMBIA2300-AarI-ccdB NLS-2×Cherry. This vector was used to create expression vectors containing two expression cassettes of genes of interest in a single T-DNA for BiFC in *M. truncatula*.

To test pCAMBIA2300-AarI-ccdB NLS-2×Cherry, the expression cassettes *STR1pro:nYFP-STR1g-t35S, STR1pro:cYFP-STR1g-t35S, STR2pro:nYFP-STR2g-t35S* and *STR2pro:cYFP-STR2g-t35S* were created in by Multi-Site Gateway recombination reactions (Table S2). *STR1pro:nYFP-STR1g-t35S* and *STR1pro:cYFP-STR1g-t35S* was amplified using primers B7952 and B7953 containing *Aar*I sites (Table S1). *STR2pro:nYFP-STR2g-t35S* and *STR2pro:cYFP-STR2g-t35S* was amplified using primers B7954 and B7955 containing *Aar*I sites (Table S1). Amplified *STR1pro:nYFP-STR1g-t35S* and *STR2pro:cYFP-STR2g-t35S or STR1pro:cYFP-STR1g-t35S* and *STR2pro:nYFP-STR2g-t35S* cassettes were used in Golden Gate cloning reaction using *Aar*I restriction enzyme (ThermoFisher Scientific) and T4 DNA Ligase (Promega) to create co-expression vectors for BiFC analysis in *M. truncatula* (Fig. S3b; Table S3).

To create co-expression vectors for BiFC analysis of 14-3-3A and CKL1/CKL2 in *M. truncatula*, the expression cassettes *14-3-3Apro:14-3-3A-nYFP-t35S, CKL1pro:CKL1g-cYFP-t35S, CKL2pro:CKL2g-nYFP-t35S* were created in Multi-Site Gateway recombination reaction (Table S2). *14-3-3Apro:14-3-3A-nYFP-t35S* was amplified using primers B7952 and B7953 containing *Aar*I sites (Table S1). *CKL1pro:CKL1g-cYFP-t35S* or *CKL2pro:CKL2g-nYFP-t35S* were amplified using primers B7954 and B7955 containing *Aar*I sites (Table S1). Amplified *14-3-3Apro:14-3-3A-nYFP-t35S* and *CKL1pro:CKL1g-cYFP-t35S* or *CKL2pro:CKL2g-nYFP-t35S* cassettes were used in Golden Gate reaction using *Aar*I restriction enzyme and T4 DNA Ligase to create co-expression vectors for BiFC analysis in *M. truncatula* (Fig. 2a; Table S3).

To create expression vectors for RNA interference of *14-3-3A* or for simultaneous RNA interference of *14-3-3A, 14-3-3B* and *14-3-3C* specifically in arbuscule containing cells the target sequences were designed using Sol Genomics Network VIGS Tool (https://vigs.solgenomics.net). The target sequence for *14-3-3A* was amplified using primers B8600 and B8601 containing *att*B1 and *att*B2 sites. Target sequences for simultaneous RNA interference were amplified using primers B8600 and B8603 (for *14-3-3A*), B8602 and B8605 (for *14-3-3B*) and B8604 and B8606 (for *14-3-3C*). DNA fragments for RNA interference of *14-3-3A, 14-3-3B* and *14-3-3C* were then assembled into a single sequence using overlapping PCR and primers B8600 and B8606 containing *att*B1 and *att*B2 sites. Obtained PCR fragments were recombined with pDONR221 to obtain pENTR L1-L2 *14-3-3A RNAi* and *14-3-3ABC RNAi* clones. These pENTR clones were used in a recombination reaction with pENTRL4-R1 *PT4pro*, containing a promoter sequence of AM specific *phosphate transporter 4* gene, and pKm42GWIWG8,1 RR destination vector (Ivanov and Harrison, 2024) using Gateway™ LR Clonase™ II Enzyme mix (ThermoFisher Scientific) to create RNAi expression vectors (Table S2).

### 7. Confocal laser-scanning microscopy and bimolecular fluorescence complementation assay

Confocal laser-scanning microscopy and bimolecular fluorescence complementation (BiFC) assays in *N. benthamiana* were performed as described previously (Ivanov & Harrison, 2024).

For confocal laser-scanning microscopy and bimolecular complementation (BiFC) assays in *M. truncatula*, the regions of transgenic roots showing fluorescence associated with fungal colonization (mCherry-BCP or NLS-2×mCherry) were excised to short pieces approximately 3-5 mm in length. The root pieces were then cut longitudinally with a double-edged razor blade (Electron Microscopy Sciences; 72000). Dissected root pieces were then placed on a glass slide in a drop of water with the cut surface facing upwards and covered by a cover slip. Roots sections were observed and fluorescence was imaged using a Leica TCS-SP5 confocal microscope (Leica Microsystems) with a 63× water-immersion objective. This imaging approach has been validated previously (Pumplin & Harrison, 2009).

For sub-cellular location, GFP was excited with the argon ion laser (488 nm; 30% power) and emitted fluorescence was collected using HyD detector from 505 to 525 nm. For BiFC assay, YFP was excited with the argon laser (514 nm; 30% power) and emitted fluorescence was collected from 529 to 549 nm. All acquisitions of YFP fluorescence for BiFC were performed at “Standard mode” and the fixed “Smart gain” value (350 V). mCherry was excited with the Diode-Pumped Solid State laser (561 nm; 30% power) and emitted fluorescence was collected using HyD detector from 605 to 630 nm. Differential interference contrast (DIC) images were collected simultaneously with the corresponding fluorescence. Images were processed using Leica LAS-AF software versions 2.6.0 (Leica Microsystems), Image J (National Institutes of Health), Adobe Photoshop 2024 version 25.1.0 and Adobe Illustrator 2024 version 28 (Adobe Systems Inc.).

### 8. Charts and Graphs

Charts and Graphs were built using PlotsOfData (https://huygens.science.uva.nl/PlotsOfData/) and edited in Adobe Illustrator 2024 version 28 (Adobe Systems Inc.).

## III. Results

### 1. A Y2H-seq screen using DMI3 and CKL2 as protein interaction baits

Yeast-2-hybrid (Y2H) analyses are widely used to identify protein-protein interactions with many variations in style and scale (Bruckner *et al*., 2009; Erffelinck *et al*., 2018). By coupling Y2H with next generation sequencing the process of handling individual clones on a clone-by-clone basis is bypassed; instead a quantitative assessment of the interactions is obtained and a rank-ordered list of candidates can be generated (Erffelinck *et al*., 2018). To enable this approach, we generated a Y2H library from cDNA of *M. truncatula* roots colonized with *Diversispora epigaea* (formerly *Glomus versiforme*). We screened the library with two bait proteins, DMI3 (Levy *et al*., 2004) and CKL2 (Ivanov & Harrison, 2024). DMI3 is a calcium calmodulin-dependent protein kinase, with a known interaction partner, INTERACTING PARTNER OF DMI3 (IPD3) (Yano *et al*., 2008; Horvath *et al*., 2011) and serves as a positive control. For each screen, the colonies from the interaction plates were pooled, the inserts were amplified and 3 replicate amplicon libraries were generated. We also generated 3 amplicon libraries from the entire Y2H library to obtain baseline information on transcript representation.

Following sequencing, an average transcript count matrix was generated for each library and the average cpm-normalized counts were ranked by abundance for each bait protein. IPD3 was the highest ranked interacting protein in the DMI3 interaction library, providing positive validation of the screening method. New candidate interacting proteins, ranked immediately below IPD3, include two S-adenosylmethionine-dependent methyltransferases, which may be interesting targets for researchers interested in DMI3 function (Table 1). By contrast, the CKL2 interaction library contained two general regulatory factor (14-3-3) proteins as the top-ranked candidates, closely followed by a third 14-3-3 protein (Table 2). A comparison to the entire library shows the relative abundance of these transcripts in the library, while comparison to the DMI3 data reveals that top-ranked candidate interactors from the CKL2 and DMI3 screens differ entirely.

**Table 2.**
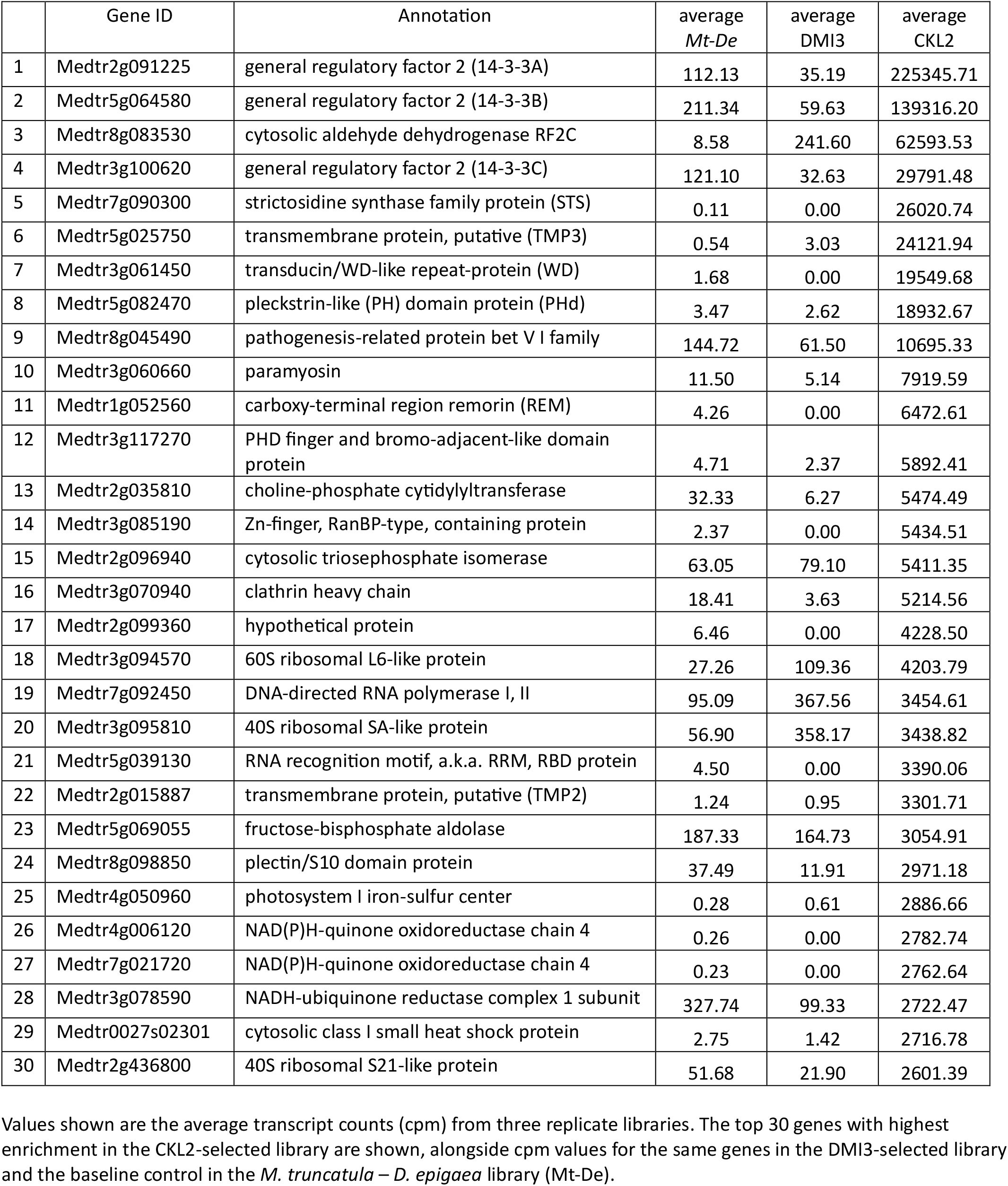
Top ranked interactors of CKL2 as identified in a Y2H-seq screen. Values shown are the average transcript counts (cpm) from three replicate libraries. The top 30 genes with highest enrichment in the CKL2-selected library are shown, alongside cpm values for the same genes in the DMI3-selected library and the baseline control in the *M. truncatula* – *D. epigaea* library (Mt-De).

### 2. Evaluation of CKL2 –protein interactions using BiFC in *Nicotiana benthamiana*

For an initial evaluation of CKL2 candidate protein interactions, we used BiFC assays (Kerppola, 2006; Kerppola, 2008) in *Nicotiana benthamiana* leaf epidermal cells. This preliminary BiFC analysis is not essential but it was useful because it allowed us to further prioritize candidate interacting proteins for subsequent analyses in *M. truncatula* mycorrhizal roots. Because there is no defined cut-off for ‘true interactors’, eight CKL2-interacting proteins were selected for evaluation in *N. benthamiana*. These included the top-ranked general regulatory factor 2 (14-3-3) proteins (*14-3-3A*, Medtr2g091225; *14-3-3B*, Medtr5g064580), a strictosidine synthase family protein (*STS*, Medtr7g090300), a putative transmembrane protein (*TMP3*, Medtr5g025750), transducin/WD-like repeat-protein (*WD*, Medtr3g061450), a pleckstrin-like (PH) domain protein (*PHd*, Medtr5g082470), a carboxy-terminal region remorin (*REM*, Medtr1g052560) and a putative transmembrane protein (*TMP2*, Medtr2g015887). Prior to the BiFC assay, we created translational GFP fusions with each gene and determined the sub-cellular location of each protein-GFP fusion in *N. benthamiana* cells (Fig. S1a; Table S2). This revealed that the 14-3-3A and 14-3-3B GFP fusion proteins located in cytoplasm and nucleoplasm; STS-GFP was detected in endoplasmic reticulum; TMP3-GFP in chloroplasts; WD-GFP in the nucleoplasm and cytoplasm; PHd-GFP on the membrane of the plant vacuole, tonoplast, and large endosomal compartments; GFP-REM on the plasma membrane; TMP2 in cytoplasm and nucleoplasm. We showed previously that CKL2-GFP is myristoylated and in *N. benthamiana* cells, it locates at the plasma membrane (Fig. S1b) (Ivanov and Harrison, 2024).

To test protein:protein interactions in BiFC assays we created translational fusions of the N-terminal half of YFP (*nYFP*) with each gene under the control of the *CaMV35S* promoter. A CKL2 translational fusion with the C-terminal half of YFP (*cYFP*) under the control of the *Arabidopsis UBQ10* promoter (*AtUBQ10pro:CKL2-cYFP*) was created and the two cassettes were then placed within a single transfer DNA (T-DNA) along with a third expression cassette (*AtUBQ10pro:GUS-mCherry)* which serves as transformation marker and allows the detection of transformed *N. benthamiana* cells (Fig. S1c; Table S3) (Ivanov and Harrison, 2024). Following *Agrobacterium tumefaciens* transformation into *N. benthamiana* leaves, transformed cells were identified by the presence of mCherry fluorescence and then imaged for YFP (Fig. 1). We detected YFP-fluorescence when *CKL2-cYFP* was co-expressed with either *14-3-3A-nYFP, 14-3-3B-nYFP, nYFP-REM* or *TMP2-nYFP* indicating a positive interaction of CKL2 with these proteins. YFP-fluorescence was not detected when *CKL2-cYFP* was co-expressed with either *STS-nYFP, TMP3-nYFP, WD-nYFP* or *PHd-nYFP* (Fig. 1). In addition, YFP-fluorescence was not detected when *CKL2-cYFP* was co-expressed with either of two negative control proteins, membrane located *nYFP-LTI6b* or cytoplasmic protein *GUS-nYFP* (Fig. 1).

**Figure 1.**
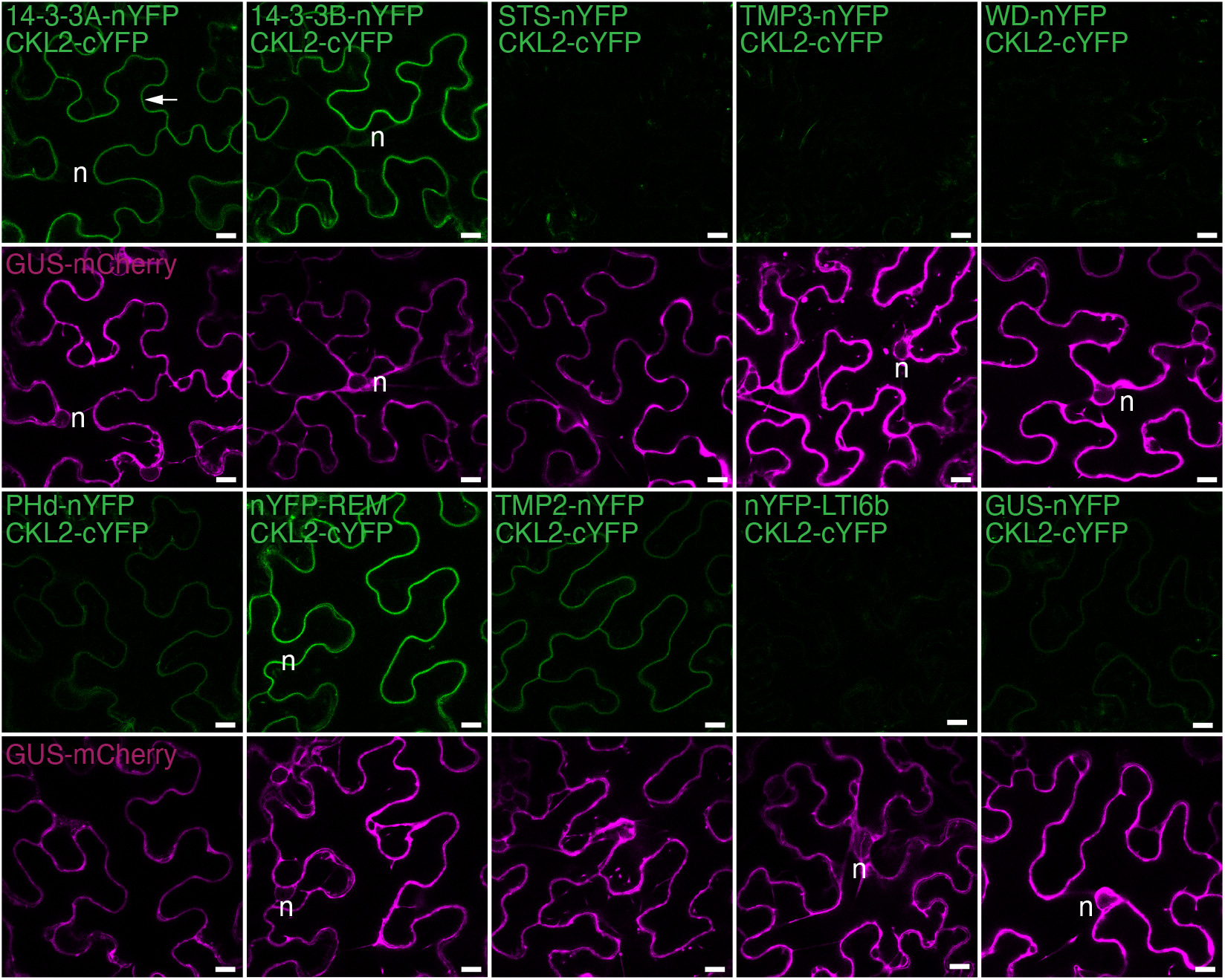
Bimolecular fluorescence complementation to assess the interaction of CKL2 and the top ranked protein interaction candidates in *N. benthamiana*. *CKL2-cYFP* was expressed (*Arabidopsis UBQ10* promoter) with top-ranked protein interaction candidates identified in the Y2H-seq screen. Protein interaction candidates were expressed (*CaMV35S promoter*) as translational *nYFP* fusions: general regulatory factors 2 (*14-3-3A* and *14-3-3B*), strictosidine synthase (*STS*), putative trans-membrane proteins (*TMP2* and *TMP3*), transducin/WD-like repeat-protein (*WD*), pleckstrin-like (PH) domain protein (*PHd*), carboxy-terminal region remorin (*REM*). Translational *nYFP* fusions of β-glucuronidase (*GUS*) and *Medicago LTI6b* were used as controls for soluble and membrane associated proteins, respectively. The reconstituted YFP fluorescence (green) was observed in leaf epidermal (pavement) cells using confocal laser-scanning microscopy. The expression and fluorescence detection of *GUS-mCherry* (magenta) was used as a marker of *Agrobacterium* infiltration and transformation. *Arrow*, plasma membrane; *n*, nucleus. Scale bar, 10 µm.

*CKL2* has a paralog, *CKL1*, which shows a slightly different subcellular location to that of CKL2; in addition to location at the plasma membrane, CKL1-GFP locates in cytoplasm and nucleoplasm (Fig. S1b) (Ivanov and Harrison, 2024). Co-expression of *CKL1-cYFP* with either *14-3-3A-nYFP* or *14-3-3B-nYFP*, resulted in a YFP signal on the plasma membrane and in the cytoplasm (Fig. S1d) but not in the nucleoplasm, even though CKL1 and both 14-3-3 proteins exist in nucleoplasm as shown by their individual GFP fusions (Fig. S1a and b) (Table S3). When *CKL1-cYFP* was co-expressed with *nYFP-REM*, the YFP-fluorescence was detected only on the plasma membrane (Fig. S1d). Thus, BiFC analysis in *Nicotiana benthamiana* supports a subset of the protein-protein interactions discovered in the Y2H-seq screen.

### 3. Generating a single T-DNA with triple gene cassette for BIFC assays in *M. truncatula* mycorrhizal roots

*N. benthamiana* is a useful heterologous system but leaf cells differ substantially from the colonized cortical cells of *M. truncatula* mycorrhizal roots. To facilitate BIFC assays in *M. truncatula* roots, where proteins interactions can be evaluated in their native location, we built a cloning system in which the vector carries gene-split YFP translational fusions under the control of their native promoters and also a transformation marker active in *M. truncatula* mycorrhizal roots. To test this triple gene cassette T-DNA, we used ABCG family transporters STR1 and STR2. The interaction of these half-size ABC transporters, at the periarbuscular membrane of colonized cortical cells, was established previously by BiFC (Zhang *et al*., 2010).

A single T-DNA construct carrying a translational fusion of *nYFP* with *STR1* under the control of the *STR1* promoter (*STR1pro:nYFP-STR1*), a translational fusion of *cYFP* and *STR2* under the control of the *STR2* promoter (*STR2pro:cYFP-STR2*) and an AM symbiosis-inducible, nuclear-located transformation marker (*BCP1pro:NLS-2×mCherry*) was generated following the scheme shown (Fig. 2a; Table S3). In addition, we created a second single T-DNA construct with reciprocal positioning of two-halves of YFP (*STR1pro:cYFP-STR1 and STR2pro:nYFP-STR2*) (Table S3). The constructs were introduced into *M. truncatula* roots via *Agrobacterium rhizogenes*-mediated transformation and the resulting composite plants, with transgenic root systems, were inoculated with the AM fungus *Rhizophagus irregularis*. For BiFC analysis, colonized regions of the root were identified based on the presence of mCherry fluorescence in the nucleus. The fluorescence from reconstituted full-length YFP was then imaged in cortical cells containing arbuscules, which is the known location of STR1 and STR2 (Fig. 2b). Consistent with our previous study (Zhang *et al*., 2010), we observed the YFP-fluorescence from the interaction of nYFP-STR1/cYFP-STR2 (cYFP-STR1/nYFP-STR2) in the PAM around arbuscule branches (Fig. 2b) (Zhang *et al*., 2010). Thus, these data indicated that the single T-DNA-triple cassette system works correctly in colonized *M. truncatula* roots.

**Figure 2.**
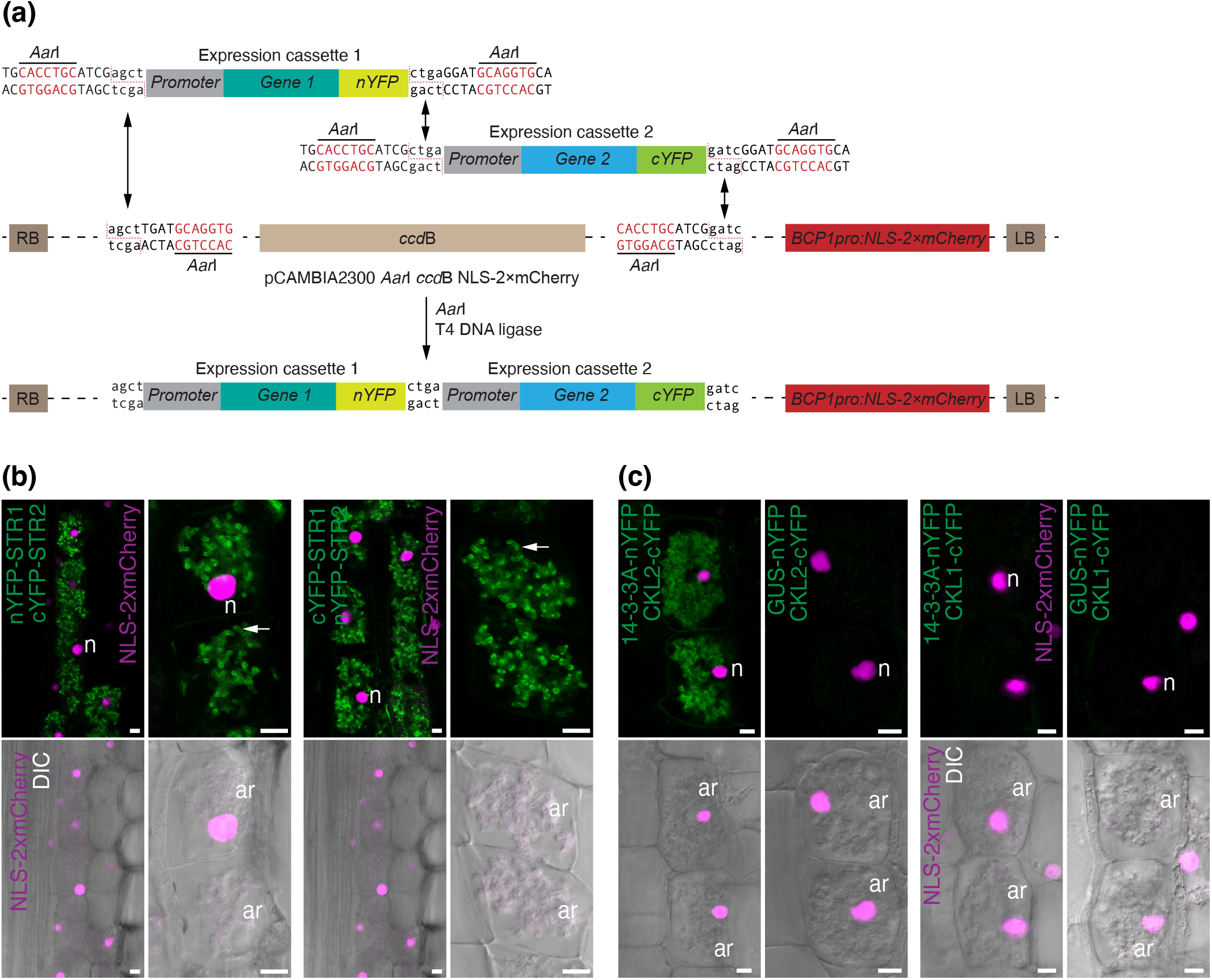
Bimolecular fluorescence complementation system to assess protein:protein interactions in *M. truncatula* root cells during AM symbiosis. **(a)** Schematic representation of the cloning approach to create binary vectors containing three expression cassettes on a single T-DNA for BiFC analysis in *Medicago* roots. Expression cassettes containing promoter and translation fusion of genes of interest with either N-terminal or C-terminal parts of YFP (*nYFP* and *cYFP*, respectively) were amplified using oligonucleotides containing recognition sites for *Aar*I restriction endonuclease. The restriction/digestion by *Aar*I was used to insert the expression cassettes into *Aar*I sites within a modified pCAMBIA2300 vector that also contains an expression cassette for a nucleus-localized fluorescent marker of intraradical AM colonization (*BCP1pro:NLS-2×mCherry*) within a single T-DNA. **(b)** As a proof of concept, two Medicago half-ABC transporters *STR* and *STR2* were co-expressed from a single T-DNA as either nYFP or cYFP translational fusions under native promoters in *Medicago* transiently transgenic roots. The reconstituted YFP fluorescence (green) was observed in root cortical cells containing arbuscules of AM fungus *Rhizophagus irregularis* using confocal laser-scanning microscopy. The nucleus-localized marker of intraradical AM colonization *NLS-2×mCherry* was expressed under AM symbiosis specific *Medicago BCP1* promoter and was used for detection of transformation and colonization in root cortical cells (magenta). The reconstituted YFP fluorescence is present at the periarbuscular membrane consistent with the published location of STR-STR2 interaction **(c)** *CKL2pro:CKL2-cYFP* or *CKL1pro:CKL1-cYFP* were co-expressed with *14-3-3Apro:14-3-3A-nYFP* from a single T-DNA in *Medicago* transiently transgenic roots. The reconstituted YFP fluorescence (green) was observed in root cortical cells containing arbuscules of AM fungus *Rhizophagus irregularis* using confocal laser-scanning microscopy. A translational fusion of nYFP with β-glucuronidase (GUS) was used as a negative control for the interaction. *Arrow*, periarbuscular membrane; *n*, nucleus; *ar*, arbuscule. DIC, differential interference contrast. Scale bar, 10 µm.

### 4. Evaluating CKL2:14-3-3A interactions in *Medicago truncatula* mycorrhizal roots

The top-ranked CKL2 interactor, 14-3-3A, was selected as a first example. Prior to evaluating the interaction of CKL2 with 14-3-3A in mycorrhizal roots, we first assessed the cellular and sub-cellular location of 14-3-3A. A translational fusion (GFP-14-3-3A) expressed from the 14-3-3A promoter revealed that 14-3-3A-GFP locates in cytoplasm and nucleoplasm of *M. truncatula* root cells, including cortical cells containing arbuscules (Fig. S2a).

To evaluate the CKL2 - 14-3-3A interaction in its native cell type and subcellular location, we created a triple gene T-DNA construct with the three components: an AM symbiosis-inducible transformation marker (*BCP1pro:NLS-2×mCherry*), a translational fusion of *nYFP* with *14-3-3A* expressed from the 14-3-3A promoter (14-3-3Apro:14-3-3A-nYFP) and a translational fusion *cYFP* with *CKL2* expressed from the CKl2 promoter (*CKL2pro:CKL2-cYFP*) (Table S3). Both the 14-3-3A and CKL2 cassettes were built from the genomic sequences to retain the native gene structure. In addition, we generated equivalent constructs with CKL1, so that CKL1-14-3-3A interactions could be assessed in mycorrhizal roots. Negative control constructs were generated with either translational fusions to GUS or a membrane protein, LTI6b (Table S3).

Following introduction into *M. truncatula* roots and inoculation with *R. irregularis*, colonized regions of the roots were identified based on the presence of mCherry fluorescence. YFP was then imaged in cortical cells containing arbuscules (Fig. 2c). Consistent with the known sub-cellular location of CKL2 (Ivanov & Harrison, 2024), roots expressing the 14-3-3A-nYFP and CKL2-cYFP construct showed a YFP signal at the PAM (Fig. 2c and Fig. S2b-c). In cortical cells with less densely branched arbuscules, the fluorescent signal from 14-3-3A-nYFP/CKL2-cYFP interaction was clearly visible at the periarbuscular membrane around arbuscular branches and trunk (Fig. S2c). A YFP signal was also visible at the plasma membrane but was considerably weaker than the signal at the PAM (Fig. 2c and Fig. S2c). We did not detect YFP in the interaction assay of 14-3-3A-nYFP/CKL1-cYFP, or in negative controls which assessed the interaction of GUS-nYFP with either CKL2-cYFP or CKL1-cYFP (Fig. 2b; Fig. S2c). Thus, the BiFC assay indicates supports the interaction of CKL2 with 14-3-3A in the native subcellular location in colonized cells of a mycorrhizal root.

### 14-3-3 proteins influence fungal colonization levels

14-3-3s are regulatory proteins without known enzymatic activity. They function as dimers, which bind to phosphorylated amino acid residues in their target proteins (Sheikh *et al*., 2024). The *A. thaliana* genome encodes several 14-3-3 isoforms, which have been named general regulatory factors (GRFs 1-14) but are also referred to as 14-3-3 proteins with subfamilies indicated by a Greek letter (Lozano-Durán & Robatzek, 2015) and divided into two subfamilies: non-epsilon, which includes GRF1 to GRF8, and epsilon, which includes GRF9 to GRF14 (Keicher *et al*., 2017). The three *Medicago* 14-3-3 proteins identified in the CKL2 Y2H-seq screen are paralogues of the *Arabidopsis* proteins as follows: Medicago 14-3-3A is a paralogue of GFR9/14-3-3µ, Medicago 14-3-3B is a paralogue of GRF1/14-3-3χ, GRF2/14-3-3ω and GRF4/14-3-3ϕ and Medicago 14-3-3C is a paralog of GRF6/14-3-3λ and GRF8/14-3-3κ.

To gain an initial evaluation of the functional relevance of 14-3-3A in AM symbiosis, we performed colonized cell-specific *14-3-3A* knock-down with a *14-3-3A* RNA interference (RNAi) construct expressed from an AM specific *phosphate transporter 4* promoter (Harrison *et al*., 2002). The *PT4pro:14-3-3A RNAi* expression cassette was introduced into *M. truncatula* roots via *Agrobacterium rhizogenes*-mediated transient transformation on a T-DNA that also contains the RedRoot marker (RedRoot; *AtUBQ10pro:DsRed*) (Ivanov & Harrison, 2024) to allow the identification of transformed roots. Following colonization, transgenic roots exhibiting DsRed fluorescence were collected and analyzed but we did not observe any effect of *14-3-3A* knock-down on AM fungal colonization when compared to control roots expressing *PT4pro:GUS RNAi* (Fig. 3a). However, within the top ten protein interaction candidates discovered in the CKL2 Y2H-seq screen, there are three 14-3-3 proteins: 14-3-3A, 14-3-3B and 14-3-3C which may suggest functional redundancy. To evaluate this, we generated a construct simultaneously targeting *14-3-3A, 14-3-3B* and *14-3-3C (PT4pro:14-3-3ABC RNAi)* and analyzed its effect on the colonization rate by AM fungus. *PT4pro:14-3-3ABC RNAi* transgenic roots showed a 31% reduction in colonization relative to control transgenic roots (Fig. 3a). Thus, these data suggest a potential role for 14-3-3-proteins in colonized cells during AM symbiosis.

**Figure 3.**
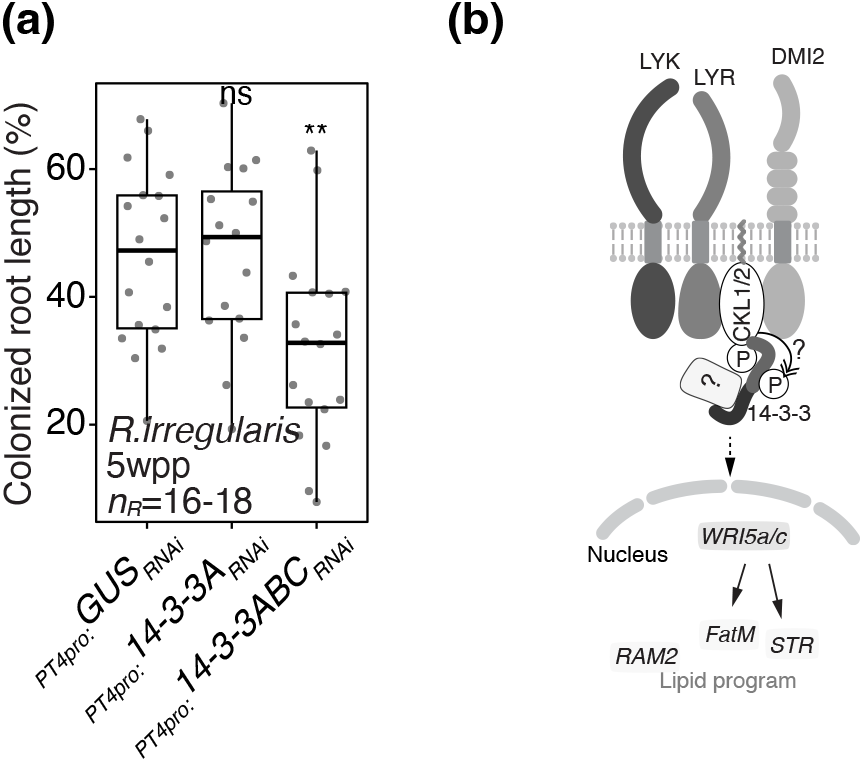
Function of 14-3-3 in AM symbiosis and proposed mode of action. **(a)** Gene knock-down of *14-3-3A, B* and *C* results in reduced colonization of Medicago transgenic roots by AM fungus *Rhizophagus irregularis* 5 weeks post planting (5 wpp). The DNA constructs for RNA interference targeting *14-3-3A* (*14-3-3A*_*RNAi*_) or *14-3-3A, B* and *C* simultaneously (*14-3-3ABC*_*RNAi*_) were expressed from AM symbiosis specific *phosphate transporter 4* promoter (*PT4 pro*) to induce gene silencing exclusively in the arbuscule containing cells. The RNA interference construct targetingβ-glucuronidase (*GUS*_*RNAi*_) was used as a control. *n*_*R*_, number of individual transgenic roots (dots); *ns*, not significant; **, p<0.01, *t*-test. **(b)** The proposed model of CKL2 and 14-3-3 interaction and function. CKL2 and its symbiotic paralogue CKL1 interact and are phosphorylated by LysM receptor-like kinases and malectin-like domain leucine-reach repeat receptor-like kinase DMI2 in cortical arbuscule containing cells of Medicago roots transducing signal to the nucleus to control lipid provisioning program regulated by transcriptional factors WRI5a and c and includes lipid biosynthesis enzymes (FATM and RAM2) and putative lipid transporters (STR), as published previously. The phosphorylated form of CKL2 (and potentially CKL1) is recognized by and interact with 14-3-3 proteins. Consequently, 14-3-3 can either facilitate the interaction of CKL2 with putative downstream signal transduction components to control lipid provisioning program or be itselfa target for phosphorylation by CKL2.

## IV. Discussion

### 1. Y2H -seq

There are several experimental approaches for identifying protein:protein interactions and the optimal approach will depend on many factors (Xing *et al*., 2016; Elhabashy *et al*., 2022). Co-immunoprecipitation or proximity-labeling methods, each coupled with mass spectrometry, are widely used and have the advantage that they identify protein interactions *in planta*. However, these approaches may be challenging if the target protein abundance is low (Qin *et al*., 2021). Additionally, in these approaches, each target protein requires tagging, unique controls and potentially optimization of protein extraction methods. Although not a panacea, Y2H screens are widely used to identify plant protein:protein interactions because they are relatively cheap and easy. However, there are disadvantages; the standard Y2H system, which involves nuclear localization of bait and prey, may not be suitable for membrane proteins. Additionally, the screens can result in false positives, so secondary evaluation *in planta* is essential. Identifying protein:protein interactions that occur exclusively in the colonized cells of mycorrhizal roots is particularly challenging and even more so for signaling proteins whose abundance may be low. A Y2H-seq screen coupled with down-stream BIFC analyses in the native protein location has several advantages. First, the screen is relatively rapid because next generation sequencing obiviates the requirement for manual purification of individual clones (Erffelinck *et al*., 2018). Additionally, multiple target proteins can be screened in a relatively short time frame and simultaneous analyses of two targets, particularly proteins with related enzymatic functions, is advantageous because candidate interactors can be compared and potential non-specific interactions flagged. Here, we screened for interactors of two kinases, DMI3 and CKL2. DMI3 served as a positive control, but in addition, the top ranked DMI3 interactors differed from those identified in the CKL2 screen, which lends support to the specificity of the interactions in both screens.

### 2. Evaluating interactions *in planta*

Prior to evaluating CKL2 interactions in *M. truncatula* mycorrhizal roots, we ran initial analyses in *N. benthamiana*. These allowed us to determine the subcellular localization of each protein and gain preliminary data about the interactions through BiFC analysis. These steps are not essential but are useful for prioritizing candidate interactors for evaluation in *M. truncatula* mycorrhizal roots. For example, several of the higher ranked CKL2 candidate interactors were localized to chloroplasts or endosomal compartments and did not show a positive interaction in the *N. benthamiana* BiFC assay. Consequently, they were not considered for analysis in *M. truncatula*.

For BiFC analysis in *M. truncatula* mycorrhizal roots, we placed the candidate interactor-split YFP fusion genes on a single T-DNA along with an AM-specific marker (*BCP1pro:NLS-2×mCherry*) in the T-DNA. This marker is expressed only in colonized regions of the roots and restriction of mCherry to the nucleus results in a strong, easily detectable signal. The marker aids in the identification of colonized cells for subsequent BIFC imaging and indicates that the cell is transformed, which is particularly important when evaluating the negative controls and cases where proteins do not interact. The use of native promoters allows evaluation of interactions under biologically relevant conditions and strengthens the conclusions. However, for proteins whose expression levels are too low for detection, stronger promoters could be utilized. Cloning schemes for generating the constructs for use in *N. benthamiana* and in *M. truncatula* are simple modular cloning cassettes generated with a combination of Gateway and Goldengate cloning technologies (Fig 2a and Fig. S1c). Negative controls should be included; optimally, these should be fusions that will show a similar sub-cellular localization as the interactors. For CKL2 interaction, we evaluated interaction with a GUS-split YFP fusion as the 14-3-3 proteins were located in the cytoplasm and nucleoplasm. Additionally, inclusion of a STR/STR2 split-YFP plasmid provides a useful point of comparison for membrane protein interactions.

### 3. CKL2 interacts with 14-3-3 proteins in mycorrhizal roots

14-3-3 proteins are regulators that bind to phosphorylated proteins and subsequently influence their functions through modification of structure and interactions. They are well known as ‘interaction hubs’ and participate in a wide array of cellular processes and signaling pathways, including modulation of membrane channels and proton pumps as well as hormone signaling and immune signaling pathways (Sheikh *et al*., 2024). In immune signaling, activation of MAP kinases by receptor-like cytoplasmic kinases, is enabled via interactions with 14-3-3 proteins (Dong *et al*., 2023).

The CKL2 Y2H screen clearly identified 14-3-3 proteins as candidate interactors and therefore these proteins became a primary focus. Focusing subsequently on 14-3-3A, we were able to demonstrate interaction with CKL2 in *M. truncatula* colonized root cells at the periarbuscular membrane, the native location of CKL2. CKL2 is also localized in the plasma membrane and based on the signal intensity from CKL2-GFP fusions, the CKL2 protein levels in the PAM and PM are similar (Ivanov and Harrison, 2024). In contrast, a CKL2-14-3-3 BIFC signal was barely detectable at the plasma membrane. As 14-3-3 proteins interact with phosphoproteins, the difference in CKL2 - 14-3-3 BIFC signal intensity at the PAM and PM may reflect differences in the CKL phosphorylation status in the PAM and plasma membranes. *In vitro* phosphorylation data showed that CKL2 can be phosphorylated by several membrane receptor kinases, so differential phosphorylation is a possibility. Subsequent interaction with 14-3-3A could provide a mechanism to direct signaling from specific membrane locations within the colonized cell.

CKL1 is located at the PAM and in the nucleus but we were unable to visualize its interaction with 14-3-3A in either location. The lack of BiFC signal may also reflect its phosphorylation status but we are cautious with this interpretation because abundance of CKL1 is considerably lower than that of CKL2. This perhaps reveals one of the caveats of this approach - that interactions with low abundance proteins may be challenging to assess via BiFC. However, continual advances in imaging sensitivity will extend the reach of this approach.

In summary, as illustrated through a case study with CKL2, we propose that Y2H-seq coupled with BIFC in *M. truncatula* mycorrhizal roots, provides a useful entry to identification of protein – protein interactions during AM symbiosis. While the consequence of the CKL2 – 14-3-3 interaction remains to be fully investigated, initial RNAi experiments suggest that the 14-3-3 proteins are required for full colonization, while also suggesting functional redundancy. The new candidate interactors identified in the ‘positive control’ DMI3 screen, may also be valid DMI3 interactors. The role of DMI3 in the nucleus and its interaction with IPD3 is well-studied, but a proportion of the DMI3 protein is localized in the cytoplasm (Fig. S3), where it may interact with other proteins. Thus, the DMI3 interaction dataset may provide a useful foundation for further investigations of DMI3.

## Acknowledgements

Funding was provided by the US National Science Foundation Plant Genome Research IOS #2139351 and by the TRIAD Foundation. The authors thank the BTI Computational Biology Center led by Dr. Suzy Strickler for RNAseq assembly and mapping. Confocal images were acquired using instrumentation in the BTI Plant Cell Imaging Center purchased with funding from NSF DBI-0618969.

## Notes

### Competing Interest Statement

The authors have declared no competing interest.

