## Supplementary materials for "Yeast-2-Hybrid-Seq and Bifluorescence Complementation Resources for assessing Protein:Protein Interactions in Arbuscular Mycorrhizal Roots: CKL2 as a Case Study"

Figure S1

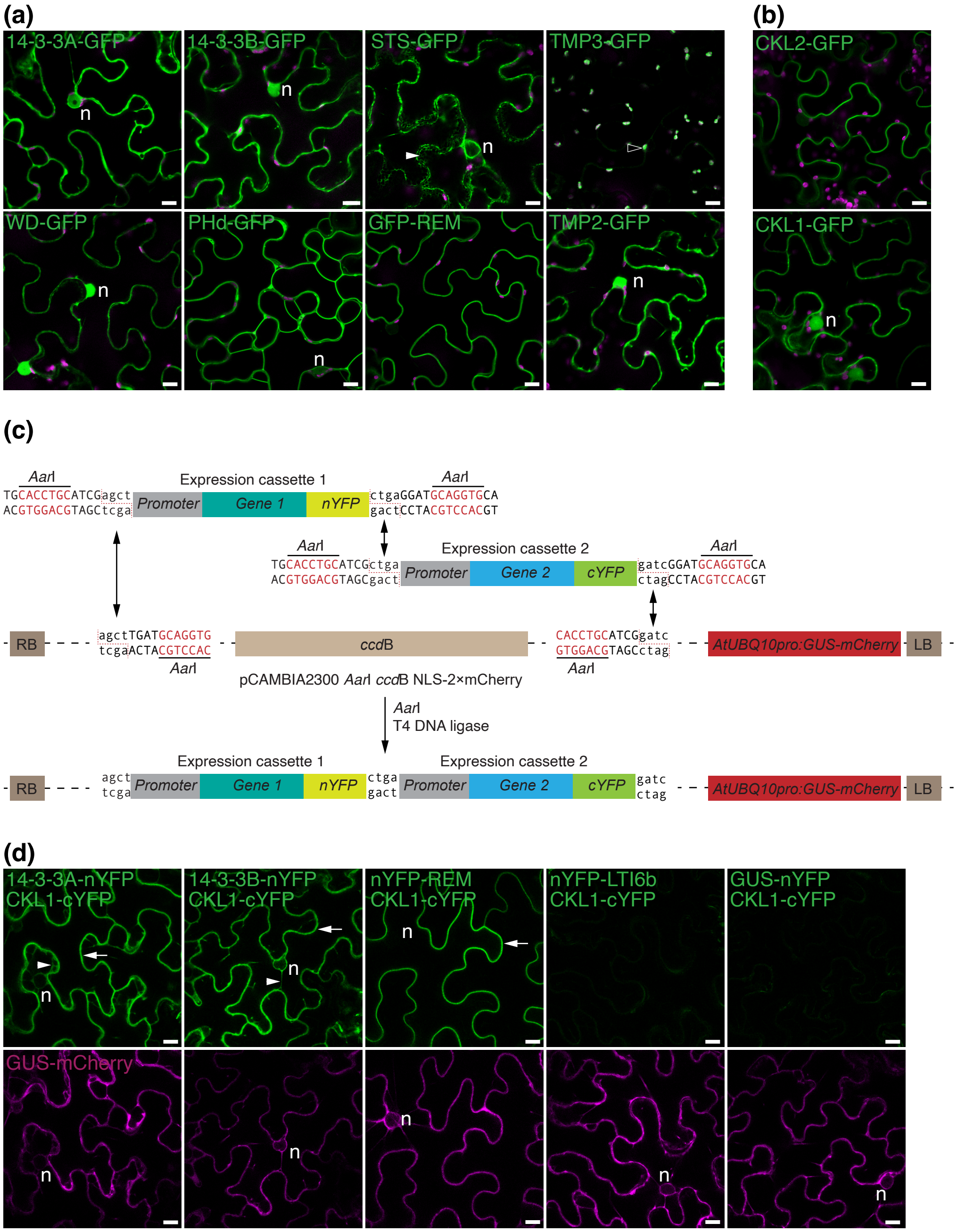
(a)
14-3-3A-GFP
14-3-3B-GFP
STS-GFP
TMP3-GFP
WD-GFP
PHd-GFP
GFP-REM
TMP2-GFP
n
(b)
CKL2-GFP
CKL1-GFP
(c)
AarI
Expression cassette 1
TGCACCTGCATCGagct
ACGTGGACGTAGCtcga
Promoter
Gene 1
nYFP
ctgaGGATGCAGGTGCA
gactCCTACGTCCACGT
Expression cassette 2
TGCACCTGCATCGctga
ACGTGGACGTAGCgact
Gene 2
cYFP
gatcGGATGCAGGTGCA
ctagCCTACGTCCACGT
RB
agctTGATGCAGGTG
tcgaACTACGTCCAC
ccdB
CACCTGCATCGgatc
GTGGACGTAGCctag
AtUBQ10pro:GUS-mCherry
LB
pCAMBIA2300 AarI ccdB NLS-2×mCherry
T4 DNA ligase
agct
tcga
ctga
gact
gatc
ctag
(d)
14-3-3A-nYFP
CKL1-cYFP
14-3-3B-nYFP
nYFP-REM
nYFP-LTI6b
GUS-nYFP
GUS-mCherry

**Figure S1. Sub-cellular location of CKL2 protein interaction candidates and their interaction with CKL1 in *N. benthamiana***

**(a)** Sub-cellular location of 14-3-3A-GFP, 14-3-3B-GFP, STS-GFP, TMP3-GFP, WD-GFP, PHd-GFP, GFP-REM and TMP2-GFP in epidermal (pavement) cells of *N. benthamiana* leaves. Confocal microscopy images. Chloroplast autofluorescence (magenta). *n*, nucleus; *arrowhead*, endoplasmic reticulum; *open arrowhead*, chloroplast. Scale bars, 10 μm. **(b)** Sub-cellular location of CKL2-GFP and CKL1-GFP in pavement cells of *N. benthamiana* leaves. Confocal microscopy images. Chloroplast autofluorescence (magenta). *n*, nucleus. Scale bars, 10 μm. **(c)** Schematic representation of the cloning approach to create binary vectors containing two expression cassettes on a single T-DNA for BiFC analysis in pavement cells of *N. benthamiana* leaves. Expression cassettes containing promoter and translation fusion of genes of interest with either N-terminal or C-terminal parts of YFP (*nYFP* and *cYFP,* respectively) were amplified using oligonucleotides containing recognition sites for *Aar*I restriction endonuclease. The restriction/digestion by *Aar*I was used to insert the expression cassettes into modified pCAMBIA2300 vector containing *Aar*I sites and an expression cassette for the cytoplasm localized fluorescent marker of *Agrobacterium* infiltration and transformation (*AtUBQ10pro:GUS-mCherry*) within a single T-DNA. **(d)** Bimolecular fluorescence complementation of CKL1 and 14-3-3A, 14-3-3B or REM in *N. benthamiana* leaves. *CKL1-cYFP* was co-expressed (*Arabidopsis UBQ10* promoter) with translational *nYFP* fusions of *14-3-3A, 14-3-3B* or *REM* (*CaMV35S promoter*) from a single T-DNA. Translational fusions of *nYFP* with β-glucuronidase (*GUS*) or with *Medicago LTI6b* were used as controls for soluble and membrane associated proteins, respectively. The reconstituted YFP fluorescence (green) was observed in leaf epidermal cells using confocal laser-scanning microscopy. The expression and fluorescence detection of *GUS-mCherry* (magenta) was used as a marker of *Agrobacterium* infiltration and transformation. *Arrow*, plasma membrane; *arrowhead*, cytoplasm; *n*, nucleus. Scale bar, 10 μm.

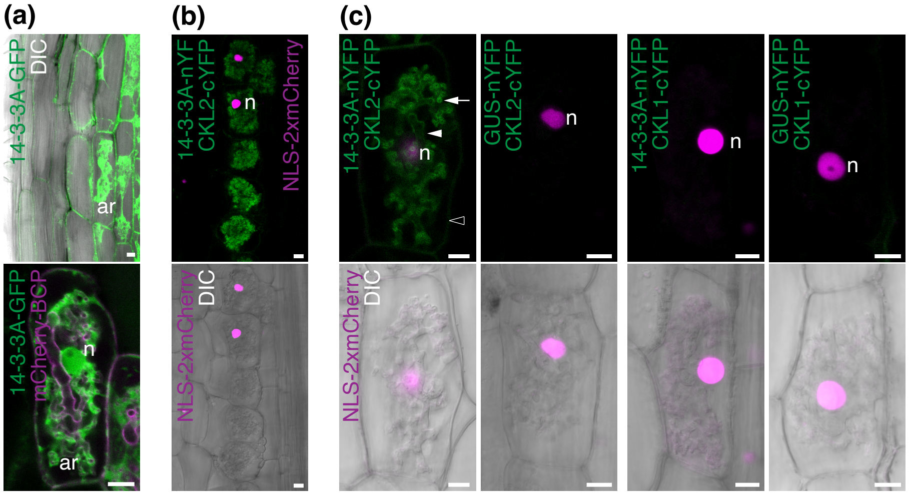


**Figure S2. Bimolecular fluorescence complementation demonstrates the interaction of CKL2-cYFP and 14-3-3A-nYFP in *M. truncatula* roots colonized with *Rhizophagus irregularis*.**

**(a)** Sub-cellular location of 14-3-3A-GFP in *Medicago* roots colonized by AM fungus *Rhizophagus irregularis*. *14-3-3Apro:14-3-3A-GFP* was co-expressed with the marker of plasma membrane and periarbuscular membrane *BCP1pro:mCherry-BCP* (magenta). The fluorescent signal from 14-3-3A-GFP (green) was detected in cytoplasm and nucleoplasm of *Medicago* root cortical cells and cortical cells containing arbuscules. *ar*, arbuscule containing cell; *n*, nucleus. DIC, differential interference contrast. Scale bar, 10μm. **(b)** and **(c)** *CKL2pro:CKL2-cYFP* or *CKL1pro:CKL1-cYFP* were co-expressed with *14-3-3Apro:14-3-3A-nYFP* from a single T-DNA in *Medicago* transgenic roots. The reconstituted YFP fluorescence (green) was observed in root cortical cells containing arbuscules of AM fungus *Rhizophagus irregularis* using confocal laser-scanning microscopy. A translational fusion of nYFP and β-glucuronidase (GUS) was used as a negative control for the interaction. A nucleus localized marker of intraradical AM colonization *NLS-2×mCherry* was expressed under an AM symbiosis-specific *Medicago BCP1* promoter and was used for detection of transformation and colonization in root cortical cells (magenta). (b) Profile of several cortical cells containing arbuscules and displaying reconstituted YFP fluorescent signal. (c) Bimolecular fluorescence complementation of CKL2-cYFP and 14-3-3A-nYFP on the periarbuscular membrane. A weak fluorescent signal was detected on the plasma membrane. *Arrow*, periarbuscular membrane of arbuscule branches; *arrowhead*, periarbuscular membrane of arbuscule trunk; *open arrowhead*, plasma membrane; *n*, nucleus; *ar*, arbuscule. DIC, differential interference contrast. Scale bar, 10μm.

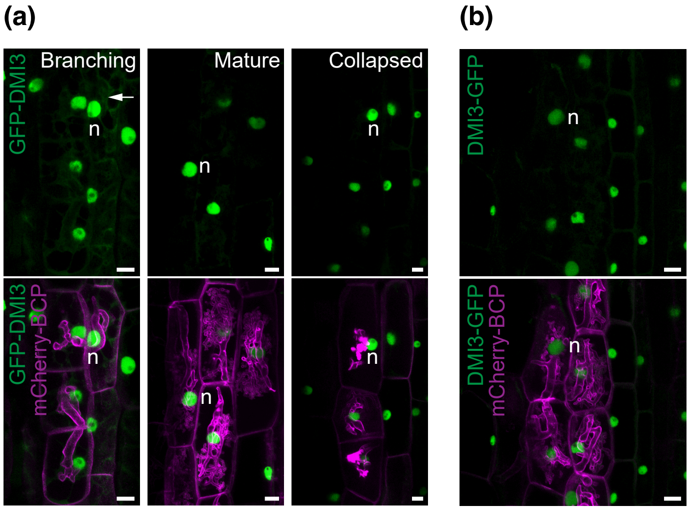


**Figure S3. Sub-cellular location of DMI3 in *M. truncatula* roots colonized with *R. irregularis***

**(a)** Sub-cellular location of GFP-DMI3 in *Medicago* roots colonized by AM fungus *Rhizophagus irregularis*. *DMI3pro:GFP-DMI3* was co-expressed with the marker of plasma membrane and periarbuscular membrane *BCP1pro:mCherry-BCP.* A fluorescent signal from GFP-DMI3 (green) was detected in the nucleus (*n*) of cortical cells containing branching, mature and collapsed arbuscules. Note, that a cytoplasmic fluorescent signal (*arrow*) is detected in cells with branching arbuscules. Confocal laser-scanning microscopy. Scale bar, 10μm. (b) Sub-cellular location of DMI3-GFP in *Medicago* roots colonized by AM fungus *Rhizophagus irregularis*. *DMI3pro:DMI3-GFP* was co-expressed with the marker of plasma membrane and periarbuscular membrane *BCP1pro:mCherry-BCP.* The fluorescent signal from DMI3-GFP (green) was detected in the nucleus (*n*) of cortical cells containing arbuscules. Confocal laser-scanning microscopy. Scale bar, 10 μm.

### Table S1. Primers used in this study

| Primer number | Primer name | Sequence |
|---|---|---|
| | | **CKL2 interactors (CDS)** |
| **B7724** | 14-3-3A-attB1-F* | **GGGGACAAGTTTGTACAAAAAAGCAGGCTTC**ATGGCTTCTTCCAAGGATCG[a] |
| **B7725** | 14-3-3A-ns-attB2-R | **GGGGACCACTTTGTACAAGAAAGCTGGGTC**CTCTGCATCGTCACCTCCAC |
| **B7881** | 14-3-3A-st-attB2-R | **GGGGACCACTTTGTACAAGAAAGCTGGGTC**TCACTCTGCATCGTCACCTC |
| **B8523** | 14-3-3B-attB1-F | **GGGGACAAGTTTGTACAAAAAAGCAGGCTTC**ATGGCAACAGCACCAACAC |
| **B8524** | 14-3-3B-st-attB2-R | **GGGGACCACTTTGTACAAGAAAGCTGGGTC**TTACTGCTCATCGGCGC |
| **B8525** | 14-3-3B-ns-attB2-R | **GGGGACCACTTTGTACAAGAAAGCTGGGTC**CTGCTCATCGGCGCC |
| **B8330** | REM-CT-attB2-F | **GGGGACAGCTTTCTTGTACAAAGTGGAA**ATGTTGAATGATCAAAGAGCTTC |
| **B8331** | REM-CT-st-attB3-R | **GGGGACAACTTTGTATAATAAAGTTGC**TTAAAAGAAGGATCTTTTGGAAGGA |
| **B7722** | TMP2-attB1-F | **GGGGACAAGTTTGTACAAAAAAGCAGGCTTC**ATGATACTAGCAACAGAAATAATAATAAC |
| **B7723** | TMP2-attB2-R | **GGGGACCACTTTGTACAAGAAAGCTGGGTC**AAAAAAACCAACCTTAAATGCA |
| **B7726** | WD-attB1-F | **GGGGACAAGTTTGTACAAAAAAGCAGGCTTC**ATGGAGGAACAATTCTCATTTAAG |
| **B7727** | WD-attB2-R | **GGGGACCACTTTGTACAAGAAAGCTGGGTC**AGCATTTTCAAAACCAAAAATTCG |
| **B7728** | TMP3-attB1-F | **GGGGACAAGTTTGTACAAAAAAGCAGGCTTC**ATGAAACTGTCTCAGGCATTGG |
| **B7729** | TMP3-attB2-R | **GGGGACCACTTTGTACAAGAAAGCTGGGTC**ATGAGCCTCGTTCAGCTTGG |
| **B7730** | PHD-attB1-F | **GGGGACAAGTTTGTACAAAAAAGCAGGCTTC**ATGGCAGCAAGTCTTTGG |
| **B7731** | PHD-attB2-R | **GGGGACCACTTTGTACAAGAAAGCTGGGTC**TTTACCATTATCATAATCAATAATTTCC |
| **B7732** | STR3-attB1-F | **GGGGACAAGTTTGTACAAAAAAGCAGGCTTC**ATGAACTCTATGATAATTGTTTCTGC |
| **B7733** | STR3-attB2-R | **GGGGACCACTTTGTACAAGAAAGCTGGGTC**TAAACTATAAACACCAATATAAGGCATTA |
| **B8607** | 14-3-3C-attb1-F | **GGGGACAAGTTTGTACAAAAAAGCAGGCTTC**ATGGGTGGTGCGATTCC |
| **B8608** | 14-3-3C-attb2-R | **GGGGACCACTTTGTACAAGAAAGCTGGGTC**AGGCTCATCTAGCTGGTCCT |
| | | **CKL2 interactors (Promoter)** |
| **B8528** | 14-3-3A-Pr-attB4-F | **GGGGACAACTTTGTATAGAAAAGTTG**GTTGGTGGCTGTGGTTCAG |
| **B8529** | 14-3-3A-Pr-attB1-R | **GGGGACTGCTTTTTTGTACAAACTTGC**CGTCGATCTCAGAGAAATTAGG |
| **B8530** | 14-3-3B-Pr-attB4-F | **GGGGACAACTTTGTATAGAAAAGTTG**GCAGTGGGTCAAGTTCAGTG |
| **B8531** | 14-3-3B-Pr-attB1-R | **GGGGACTGCTTTTTTGTACAAACTTGC**GGATGAATCTGGTTGAAGGAA |
| | | **14-3-3 RNAi** |
| **B8600** | 14-3-3A-Ri-attB1-F | **GGGGACAAGTTTGTACAAAAAAGCAGGCTTC**AGATGGTGGATTCAATGAAGAA |
| **B8601** | 14-3-3A-Ri-attB2-R | **GGGGACCACTTTGTACAAGAAAGCTGGGTC**AGTAAAACACAGTTGATTCACCA |
| **B8602** | 14-3-3AB-F | CAACTGTGTTTTACTTGGCAACAGCACCAA |
| **B8603** | 14-3-3AB-R | TTGGTGCTGTTGCCAAGTAAAACACAGTTG |
| **B8604** | 14-3-3CB-F | AAGCTGAACTTTCCATGGGTGGTGCGATTC |
| **B8605** | 14-3-3CB-R | GAATCGCACCACCCATGGAAAGTTCAGCTT |
| **B8606** | 14-3-3C-Ri-attB2 | **GGGGACCACTTTGTACAAGAAAGCTGGGTC**ATTCAACTTTGGATCTGTAATCCTT |
| | | **Yeast-2-hybrid screen** |
| **B5940** | | AATATACCTCTATACTTTAACGTC |
| **B2008** | | GAAGTGTCAACAACGTATCTACC |
| | | **STR1 and STR2** |
| **B3523** | pMtSTR-attB4 F | **GGGGACAACTTTGTATAGAAAAGTTG**CTTTGATTTATATCCCCGTAGTGG |
| **B3524** | pMtSTR-attB1 R | **GGGGACTGCTTTTTTGTACAAACTTGC**GCGTAGTAGAGCAATCACTTCTTATG |
| **B7303** | STR2-Pr-attB4-F | **GGGGACAACTTTGTATAGAAAAGTTG**GTAGTGCATAAGATGAAACACTGGA |
| **B7304** | STR2-Pr-attB1-R | **GGGGACTGCTTTTTTGTACAAACTTGC**TTTCTTGAGGTGCAAGTTAAGTTTT |
| **B3913** | STR attB2 Fw | **GGGGACAGCTTTCTTGTACAAAGTGGAA**ATGGCAAGGCTCGAGAGG |
| **B3914** | STR attB3 Rv | **GGGGACAACTTTGTATAATAAAGTTGC**TTTTCTTTCATTTTTGGAGTAG |
| **B3915** | STR2 attB2 F | **GGGGACAGCTTTCTTGTACAAAGTGGAA**ATGAAAACACAAGGTCTTGAACTTG |
| **B3916** | STR2 attB3 R | **GGGGACAACTTTGTATAATAAAGTTGC**GGACCTTTGATTTTTTGATGCAAAACG |
| | | **pCAMBIA2300-AarI-ccdB-NLS-×2mCherry** |
| **B3189** | NLS-attB1-F | **GGGGACAAGTTTGTACAAAAAAGCAGGCTTC**ATGCAGCCTTCTCTTAAACG |
| **B3037** | mCherry ns-attB2 | **GGGGACCACTTTGTACAAGAAAGCTGGGTC**CTTGTACAGCTCGTCCATGC |
| **B3901** | BCPp-KpnI-F | GG**GGTACC**GAGAGAGGGAGATGTGTTTTTAAGG[b] |
| **B4704** | T35S-SacI-R | GG**GAGCTC**GTCACTGGATTTTGGTTTTAGGA |
| **B7952** | AarI-F1 | TG**cacctgc**ATCGAGCTTACGCCAAGCTATCAACTTTGT[c] |
| **B7953** | AarI-R1 | TG**cacctgc**ATCCTCAGCACTGGATTTTGGTTTTAGGAA |
| **B7954** | AarI-F2 | TG**cacctgc**ATCGCTGATACGCCAAGCTATCAACTTTGT |
| **B7955** | AarI-R2 | TG**cacctgc**ATCCGATCCACTGGATTTTGGTTTTAGGAA |

*, *F* – forward, *R* – revers

a, in Bold are *att*B sites

b, in Bold and Underlined are sites for restriction endonucleases

c, in Bold and lowercase are *Aar*I sites

ns, no stop codon

st, stop

p or Pr, promoter

### Table S2. Expression vectors used in this study

| pENTR L4-R1 | pENTR L1-L2 | pENTR R2-L3 | Destination vector (R4-R3) |
|---|---|---|---|
| | **CKL2 interactors** | | |
| *CaMV35Spro* | *14-3-3Ac-ns* | *GFP* | pK7m34GW |
| *CaMV35Spro* | *14-3-3Bc-ns* | *GFP* | pK7m34GW |
| *CaMV35Spro* | *STR3c-ns* | *GFP* | pK7m34GW |
| *CaMV35Spro* | *TMP3c-ns* | *GFP* | pK7m34GW |
| *CaMV35Spro* | *WDc-ns* | *GFP* | pK7m34GW |
| *CaMV35Spro* | *PHdc-ns* | *GFP* | pK7m34GW |
| *CaMV35Spro* | *GFP* | *REM-CT* | pK7m34GW |
| *CaMV35Spro* | *TMP2c-ns* | *GFP* | pK7m34GW |
| *CaMV35Spro* | *GUS-ns* | *GFP* | pK7m34GW |
| *CaMV35Spro* | *14-3-3Ac-ns* | *nYFP* | pK7m34GW |
| *CaMV35Spro* | *14-3-3Bc-ns* | *nYFP* | pK7m34GW |
| *CaMV35Spro* | *STR3c-ns* | *nYFP* | pK7m34GW |
| *CaMV35Spro* | *TMP3c-ns* | *nYFP* | pK7m34GW |
| *CaMV35Spro* | *WDc-ns* | *nYFP* | pK7m34GW |
| *CaMV35Spro* | *PHdc-ns* | *nYFP* | pK7m34GW |
| *CaMV35Spro* | *nYFP* | *REM-CT* | pK7m34GW |
| *CaMV35Spro* | *TMP2c-ns* | *nYFP* | pK7m34GW |
| *CaMV35Spro* | *GUS-ns* | *nYFP* | pK7m34GW |
| *MtBCP1pro* | *NLS-mCherry* | *mCherry* | pK7m34GW |
| *14-3-3Apro* | *14-3-3Ag* | *GFP* | pK7m34GW-BCP-R |
| *14-3-3Apro* | *14-3-3Ag* | *nYFP* | pK7m34GW |
| *CKL1pro* | *CKL1g-ns* | *cYFP* | pK7m34GW-BCP-R |
| *CKL1pro* | *CKL2g-ns* | *cYFP* | pK7m34GW-BCP-R |
| *AtUBQ10pro* | *GUS* | *nYFP* | pK7m34GW |
| *PT4pro* | *14-3-3A RNAi* | | pKm42GWIWG8,1 RR |
| *PT4pro* | *14-3-3ABC RNAi* | | pKm42GWIWG8,1 RR |
| | **STR1 and STR2** | | |
| *STR1pro* | *GFP* | *STR1g* | pK7m34GW-BCP-R |
| *STR2pro* | *GFP* | *STR2g* | pK7m34GW-BCP-R |
| *STR1pro* | *nYFP* | *STR1g* | pK7m34GW-BCP-R |
| *STR1pro* | *cYFP* | *STR1g* | pK7m34GW-BCP-R |
| *STR2pro* | *nYFP* | *STR2g* | pK7m34GW-BCP-R |
| *STR2pro* | *cYFP* | *STR2g* | pK7m34GW-BCP-R |
| | **DMI3** | | |
| *DMI3pro* | *DMI3c* | *GFP* | pK7m34GW-BCP-R |
| *DMI3pro* | *GFP* | *DMI3c* | pK7m34GW-BCP-R |

* Abbreviations used in the Table S2:
  - pro, promoter
  - g, genomic sequence
  - c, coding sequence
  - ns, no stop codon
  - st, stop codon

### Table S3. Vectors for co-expression analysis (BiFC) used in this study

| Backbone | *Expression cassette 1* | *Expression cassette 1* |
|---|---|---|
| | ***For N. benthamiana*** | |
| pCAMBIA3300-AarI-ccdB GmC | *CaMV35Spro:14-3-3Ac-nYFP-35St* | *AtUBQ10pro:CKL2-cYFP-35St* |
| pCAMBIA3300-AarI-ccdB GmC | *CaMV35Spro:14-3-3Bc-nYFP-35St* | *AtUBQ10pro:CKL2-cYFP-35St* |
| pCAMBIA3300-AarI-ccdB GmC | *CaMV35Spro:STR3c-nYFP-35St* | *AtUBQ10pro:CKL2-cYFP-35St* |
| pCAMBIA3300-AarI-ccdB GmC | *CaMV35Spro:TMP3c-nYFP-35St* | *AtUBQ10pro:CKL2-cYFP-35St* |
| pCAMBIA3300-AarI-ccdB GmC | *CaMV35Spro:WDc-nYFP-35St* | *AtUBQ10pro:CKL2-cYFP-35St* |
| pCAMBIA3300-AarI-ccdB GmC | *CaMV35Spro:PHdc-nYFP-35St* | *AtUBQ10pro:CKL2-cYFP-35St* |
| pCAMBIA3300-AarI-ccdB GmC | *CaMV35Spro:nYFP-REM-CTc-35St* | *AtUBQ10pro:CKL2-cYFP-35St* |
| pCAMBIA3300-AarI-ccdB GmC | *CaMV35Spro:TMP2c-nYFP-35St* | *AtUBQ10pro:CKL2-cYFP-35St* |
| pCAMBIA3300-AarI-ccdB GmC | *CaMV35Spro:GUS-nYFP-35St* | *AtUBQ10pro:CKL2-cYFP-35St* |
| pCAMBIA3300-AarI-ccdB GmC | *CaMV35Spro:14-3-3Ac-nYFP-35St* | *AtUBQ10pro:CKL1-cYFP-35St* |
| pCAMBIA3300-AarI-ccdB GmC | *CaMV35Spro:14-3-3Bc-nYFP-35St* | *AtUBQ10pro:CKL1-cYFP-35St* |
| pCAMBIA3300-AarI-ccdB | *CaMV35Spro:14-3-3Ac-GFP-35St* | *AtUBQ10pro:CKL1-3×HA -35St* |
| pCAMBIA3300-AarI-ccdB | *CaMV35Spro:14-3-3Ac-GFP-35St* | *AtUBQ10pro:CKL2-3×HA -35St* |
| pCAMBIA3300-AarI-ccdB | *CaMV35Spro:GUS-GFP-35St* | *AtUBQ10pro:CKL1-3×HA -35St* |
| pCAMBIA3300-AarI-ccdB | *CaMV35Spro:GUS-GFP-35St* | *AtUBQ10pro:CKL2-3×HA -35St* |
| | ***For M. truncatula*** | |
| pCAMBIA3300-AarI-ccdB NLS-2×mCherry | *STR1pro:nYFP-STR1g-35St* | *STR2pro:cYFP-STR2-35St* |
| pCAMBIA3300-AarI-ccdB NLS-2×mCherry | *STR1pro:cYFP-STR1g-35St* | *STR2pro:nYFP-STR2-35St* |
| pCAMBIA3300-AarI-ccdB NLS-2×mCherry | *14-3-3Apro:14-3-3Ag-nYFP-35St* | *CKL1pro:CKL1-cYFP-35St* |
| pCAMBIA3300-AarI-ccdB NLS-2×mCherry | *14-3-3A pro:14-3-3Ag-nYFP-35St* | *CKL2pro:CKL2-cYFP-35St* |
| pCAMBIA3300-AarI-ccdB NLS-2×mCherry | *AtUBQ10pro:GUS-nYFP-35St* | *CKL1pro:CKL1-cYFP-35St* |
| pCAMBIA3300-AarI-ccdB NLS-2×mCherry | *AtUBQ10pro:GUS-nYFP-35St* | *CKL2pro:CKL2-cYFP-35St* |

* Abbreviations used in the Table S2:
    pro, promoter
    g, genomic sequence
    c, coding sequence
    ns, no stop codon
    st, stop codon
    t, terminator
